# Challenges in Brillouin microscopy: Addressing heterogeneity and contamination for reliable biomechanical mapping of fresh brain tumors

**DOI:** 10.64898/2026.04.17.719181

**Authors:** Ortrud Uckermann, Tina Leonidou, Jan Rix, Achim Temme, Ilker Y. Eyüpoglu, Roberta Galli

## Abstract

**Objective and Rationale:** Brain biomechanics is a rapidly evolving field, with mechanical properties influencing both normal development and pathological conditions such as cancer. Brillouin microscopy, a non-contact optical technique, offers a promising approach for studying the biomechanics of fresh brain tumors and organoids at subcellular resolution. However, challenges such as tissue heterogeneity and signal attenuation necessitate an in-depth evaluation of measurement strategies and potential confounding factors.

**Methods:** Fresh human brain tumor samples and tumor organoids were analyzed using Brillouin microscopy with 780 nm excitation. Measurements in the form of maps of various size were performed, and the impact of focal position, tissue heterogeneity and blood contamination on Brillouin data was assessed. Complementary Raman spectroscopy was performed as reference for tissue composition.

**Results:** Brillouin signal intensity decreased exponentially with depth, with valid measurements achievable up to 80 µm. Low signal intensities at greater depths compromised data reliability due to fitting algorithm limitations. Structural heterogeneity, including different cell types, differentially affected signal attenuation. Blood contamination was identified as a major confounder, leading to erroneous biomechanical readings. Brillouin intensity maps provided essential quality control for accurate data interpretation. Raman spectroscopy identified the presence of blood and tissue-specific biochemical signatures, reinforcing the importance of multimodal analysis.

**Conclusions:** Brillouin microscopy can effectively probe biomechanical properties of fresh brain tumors but is influenced by tissue heterogeneity and contaminants. Proper sample preparation, strategic focal positioning, and complementary techniques like Raman spectroscopy are critical for ensuring reliable data. These findings contribute to refining Brillouin microscopy protocols for neuro-oncological research and potential future clinical applications.

## 1. Introduction

The field of brain biomechanics is gaining increasing scientific attention. Mechanical properties and signaling, in addition to the well-known chemical and electrical signals, are important parameters influencing brain development. Furthermore, the biomechanics of cells and tissues play a role in pathological conditions, including cancer [1].

Nervous tissue is composed of different cell types (various types of neurons and glia cells), extracellular matrix and blood vessels and, therefore, the brain structure is heterogeneous. The distribution of cell bodies varies greatly between different regions of the brain. Functional nuclei and certain layers of specific brain structures such as the hippocampus or cerebellum are characterized by high cell densities, while fiber tracts consist mainly of myelinated axons.

Given its structural heterogeneity, it is not surprising that the brain has regionally different biomechanical properties. Cell bodies of freshly isolated pyramidal neurons and astrocytes displayed elastic moduli of 1 kPa and 0.7 kPa, respectively, and are very soft compared to other cell types [2]. They have heterogeneous viscoelastic properties due to presence of cell organelles [2]. The stiffest component of the brain are myelinated axons, with Young’s modulus ∼12 kPa [3]. Mechanical tests on fresh tissue samples in comparison to immunohistochemical analysis showed that the mechanical signature of tissue does not result from only one tissue component but is influenced by several tissue constituents for mouse [4] and human brain [5]. Magnetic resonance electrography (MRE) confirmed regional differences in mechanical properties in the healthy human brain in vivo [6].

Similarly, human brain tumors exhibit great heterogeneity regarding biomechanical properties and are considered to be softer than surrounding healthy tissue. This was shown by MRE in glioblastoma patients in vivo [7,8] and mechanical properties were confirmed to differ between glioblastoma and peritumoral regions by indentation measurements on fresh biopsies [9]. Optical coherence elastography (OCE) [10] enables to retrieve elastographic maps of displacement, stiffness and Young’s modulus that can be overlaid with optical coherence tomography (OCT) images showing the brain tumor morphology [11,12]. Atomic force microscopy (AFM) provides information on Young’s modulus and allows drawing conclusions on tissue and cell biomechanics of the brain [13]. It was applied to analyze fresh human brain tumor and non-tumor samples from a few patients, and an increase in Young’s modulus was observed for glioblastoma, however with a large range [14]. AFM measurements showed highly heterogeneous values for Young’s modulus in pituitary gland tissue [15]. Cellular stiffness has been suggested to influence the biomechanical properties of whole tissue by comparing results of MRE and probing of isolated cells using an optical stretcher [16], underscoring the importance of combining different methods for a comprehensive understanding of the role of biomechanics in cancer. Different methods operate at different scales ranging from whole tissue to the subcellular level and probe physically different aspects of biomechanics. Technical limitations and constraints of the different methodologies must be considered when interpreting the data.

Recent technical developments and research have established Brillouin spectroscopy as a promising alternative method to study biomechanics, allowing investigations at subcellular resolution [10,17]. It has the advantage of being a non-contact, optical approach and no direct mechanical action is applied to the sample during measurements. Recent reviews highlight the theoretical and technical aspects and showcases examples of applications on biological samples [18,19]. Cellular and subcellular biomechanical properties have been studied in a variety of cell types [20,21]. So far, Brillouin microscopy has mostly been applied to transparent specimens and is ideally suited to study cornea, and may also serve as a diagnostic tool [22]. Studies of healthy and pathological bladder tissue [23] and on retina [24] confirm that Brillouin microscopy has the potential to provide relevant data also when applied to turbid, cell-rich samples.

Brillouin microscopy of fresh mouse brain tissue highlighted differences in Brillouin shift between different brain regions, but did not address different nervous tissue compositions [25]. Other studies have investigated formalin-fixed mouse brain tissue [26,27]. A recent preprint describes Brillouin mapping of zebrafish larvae brains, whose brain composition is not comparable with the one of mammals [28]. Only sporadic studies on in vitro glioblastoma models are available, addressing subcellular differences [29], cell stiffness in different environments [30,31] and effects of temozolomide [32]. Unlike transparent or homogeneous biological tissues (e.g., cornea, embryo), brain tissue poses specific challenges such as: substantial depth-dependent signal degradation due to significant scattering and absorption by densely packed, irregularly organized cellular and extracellular components, high lipid content of myelinated axons in white matter and presence of extra- and intracellular lipid droplets especially in brain tumors. Moreover, potential blood contamination can obscure biomechanical signatures in brain tumor samples.

To advance neurooncological research in biomechanics using Brillouin microscopy, it is necessary to study patient-derived clinical specimens. Given the anticipated substantial heterogeneity in biomechanics, it is necessary to understand whether local variations in Brillouin parameters genuinely reflect alterations or are merely artefacts. Therefore, it is essential to establish measurement strategies and protocols, while also identifying and addressing potential sources of error and limitations. Here, we used fresh tissue samples from human brain tumors and tumor organoids. First, we investigated the influence of the focal position on the quality of the Brillouin signal, taking into account the three-dimensional tissue structure and the differential influence of cellular structures. Moreover, the potential confounding effect of blood was examined.

## 2. Materials and Methods

### Preparation of tissue slices

Brain tumor samples were obtained from routine brain tumor surgeries and were transferred to artificial cerebrospinal fluid (ACSF, containing 129 mM NaCl, 3 mM KCl, 1 mM CaCl_2_, 10 mM HEPES, 0.2 mM MgCl_2_, 20 mM Glucose) immediately after resection. The tissue sample was embedded in low melting point agarose (5%, Plaque Agarose, Biozym Scientific GmbH, Hessisch Oldendorf, Germany), slices of 500 µm thickness were prepared using a vibratome (VT1200S, Leica Biosystems Nussloch GmbH, Nussloch, Germany) and transferred to ACSF solution for Brillouin microscopy.

### Preparation of glioblastoma organoids

Brain tumor organoids were prepared of a fresh brain tumor sample of a glioblastoma (GBM) patient according to an established protocol [33] and analyzed after five weeks. All media and supplements were obtained from Thermo Fisher Scientific, Schwerte, Germany unless otherwise stated. The resected tissue sample was transferred Hibernate A medium at 4 °C. Afterwards it was placed in a sterile Petri dish with H+GPSA medium (Hibernate A, 1× GlutaMax, 1× PenStrep, and 1× Amphotericin B) and cut into small pieces (0.5 to 1 mm) using two scalpels. Subsequently, tissue pieces were washed with H+GPSA Medium and Hank’s Balanced Salt Solution (HBSS), briefly vortexed with 1× RBC Lysis Buffer (Red Blood Cell Lysis Solution 10×, Miltenyi Biotech, Bergisch Gladbach, Germany 130-094-183), and incubated at room temperature. After centrifugation (300g, 5 min at room temperature) the supernatant was carefully removed. The tissue pieces were transferred to ultra-low attachment 6-well plates with 4 ml GBO medium (50% DMEM:F12, 50 % Neurobasal, 1× GlutaMax, 1× NEAAs, 1× PenStrep, 1× N2 supplement, 1× B27 w/o vitamin A supplement, 55 µM 2mercaptoethanol (Merck KGaA, Darmstadt, Germany), and 2.5 μg/ml human insulin (Merck KGaA) per well. Cultures were maintained at 37 °C with 5% CO_2_, on a shaking platform set at 70 rpm, with a 75% medium change every 48 hours. After two weeks, the organoids were cut into smaller pieces ∼ 200 – 500 μm and transferred to 96-well U-bottom plates with 200 μL of GBO medium.

### Combined Brillouin - Raman system

Brillouin spectra were acquired using a custom build system [29], additionally allowing simultaneous Raman spectroscopic measurements (Figure 1). The system is comprised of five functional blocks:

**Figure 1.**
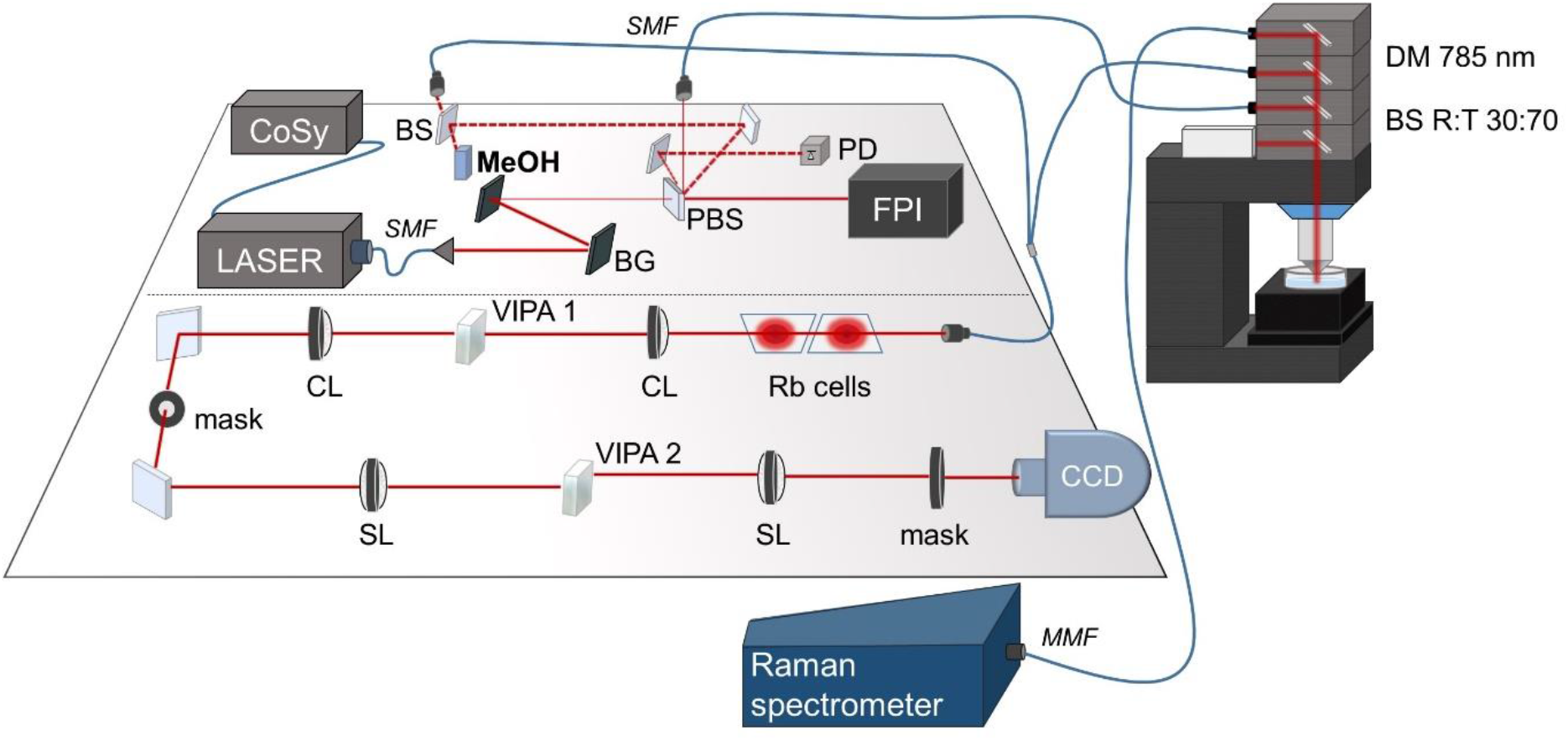
Schematic illustration of the Brillouin-Raman system. CoSy: compact saturation spectroscopy module, SMF: single mode fiber, MMF: multimode fiber, BG: Bragg grating, PBS: polarizing beam splitter, FPI: Fabry–Pérot interferometer, PD: photodiode, BS: neutral beam splitter, DM: dichroic mirror, Rb cells: rubidium cells, CL: cylindrical lens (f = 200 mm), SL: spherical lens (f = 200 mm), VIPA: virtually imaged phased array, CCD charged coupled device. Solid lines represent the main beam path, while dashed lines illustrate ghost beams arising from the PBS.

#### 1) Excitation

Narrow band laser excitation is provided by a TApro diode laser tuned at the wavelength of 780.24 nm by a CoSy saturation spectroscopy module (both TOPTICA Photonics AG, Gräfelfing, Germany). Then, the amplified spontaneous emission is removed using two Bragg gratings (NoiseBlockTM, ONDAX Inc, Monrovia, United States) and a Fabry–Pérot interferometer (LightMachinery Inc., Nepean, Canada), reaching a background suppression of – 90 dB in the spectrometer. The role of the FPI is to suppress the laser’s background originating from amplified spontaneous emission. It is locked to the maximum laser light transmission (by measuring the intensity of a ghost beam with the photodiode).

#### 2) Microscop

The laser beam is propagated to the microscope (WITec alpha 300R; WITec GmbH, Ulm, Germany), where it passes a beam splitter with reflection:transmission ratio of 30:70 and is focused on the sample using a 40× water-dipping objective (Zeiss N-Achroplan 40×/0.75 NA). The lateral optical resolution of the system was 0.6 µm (FWHM of the laser focus). The incident power on the samples was 20 mW, as measured at the sample plane using a laser power meter. The backscattered light is separated by a dichroic mirror with edge at 785 nm and directed to either the Brillouin or Raman spectrometer.

#### 3) Reference arm

A ghost beam of the polarizing beam-splitter is directed to a cuvette containing methanol and the scattered light is coupled into the Brillouin spectrometer.

#### 4) Brillouin spectrometer

Removal of Rayleigh scattering is obtained by two rubidium vapor cells (TG-ABRB-Q, Precision Glassblowing Inc, Englewood, United States) heated to 50°C. Then, the light passes two perpendicularly oriented virtually imaged phased arrays (VIPAs) having different free spectral ranges (FSR1 = 15 GHz and FSR2 = 21.6 GHz, OP-6721-6743-4, OP-6721-4720-4, LightMachinery Inc., Nepean, Canada), resulting in a rectangular interference pattern that is focused via a magnification objective (InfiniProbe TS-160; Infinity Photo-Optical Company, Centennial, CO) on a CCD camera (iDUS 420A-BR DD, Andor Technology Ltd., Belfast, Northern Ireland). The resulting spectral resolution was 44 MHz/pixel.

#### 5) Raman spectrometer

The Raman scattered light is coupled into a commercial Raman spectrometer (UHTS 400, WITec GmbH, Ulm, Germany) equipped with an optical grating with 300 grooves/mm for spectral coverage at 3 cm^−1^ resolution and CCD camera (Andor iDUS 401A-BR-DD-352, Andor Technology Ltd, Belfast, Northern Ireland).

### Data acquisition

Acquisition of Brillouin and Raman spectra were performed using WITec Suite FIVE (WITec GmbH, Ulm, Germany). Brillouin data sets were acquired in the form of maps with 1 µm or 2 µm step size and integration time of 0.2 s per point, as indicated in the figure captions. Combined Brillouin and Raman spectral measurements were performed with 30 s integration time and four accumulations.

### Data processing and analysis

Data was processed using MATLAB2023a (MathWorks Inc., Natick, MA). Lorentzian curve fitting based on MATLAB function lsqnonlin was used to extract the Brillouin peak parameters, taking into account the known methanol Brillouin shift of ν_B_ = 3.81 GHz and the fact that Stokes and anti-Stokes signals have equal absolute frequencies [29]. The frequency shift ν_B_ (center), the intensity (maximum) and the linewidth Γ_B_ (full width at half maximum) of the Brillouin band were extracted and displayed as color coded maps. All reported Γ_B_ are the measured values obtained from the Lorentzian fitting. The system broadening with the used microscope objective amounts to 0.33 GHz (FWHM of the Rayleigh band). Normalization of Brillouin intensity was performed by calculation of relative values to reference intensity measured outside the tissue in ACSF.

Raman spectra were baselined and intensity corrected using the MATLAB functions msbackadj and msnorm. Prism 10 (GraphPad Software) was used to plot graphs.

## 3. Results

Fresh tissue samples are complex, three-dimensional objects which consist of different types of cells, blood vessels and extracellular matrix. Since Brillouin spectroscopy is an optical method, the tissue properties might affect the properties of the Brillouin signal, the analysis and thus ultimately also the interpretation of the biomechanical properties through scattering, absorption and presence of different cellular components.

As a starting point, we investigated the changes of the Brillouin signal in different tissue depths. Figure 2A shows maps of Brillouin shift, intensity and bandwidth of the same xy-position of a fresh tissue sample (brain metastases of squamous cell carcinoma), which were acquired along the z-axis starting above the sample’s surface at z = 80 µm and moving down to z = -160 µm inside the sample. The Figure 2B and 2C show the median ν_B_, Γ_B_ and the median Brillouin intensity for each map, respectively. The maps above the tissue’s surface (z = 80 to 20 µm) probed the ACSF and show a uniform ν_B_ ∼ 5.16 GHz in the entire map. On the surface of the sample (z = 0), as identified in the bright field image (Figure 2D), a median ν_B_ = 5.43 GHz and Γ_B_ = 0.76 are observed, consistent with previous data on nervous tissue samples acquired at the same excitation wavelength [34]. Hence, tissue structures can be recognized by different values for ν_B_ and Γ_B_. Concurrently, the Brillouin intensity undergoes a reduction upon entering the tissue (in median a reduction to 0.69 at z = 0 compared to the intensity in medium above the sample, Figure 2B). Within the tissue, the Brillouin intensity further decreases with increasing measurement depth. An exponential fitting of the median intensity values (blue line in Fig. 2C) retrieved an attenuation length of 63 µm. For z = -20 to -80 µm, median ν_B_ between 5.4 and 5.5 GHz are observed.

**Figure 2:**
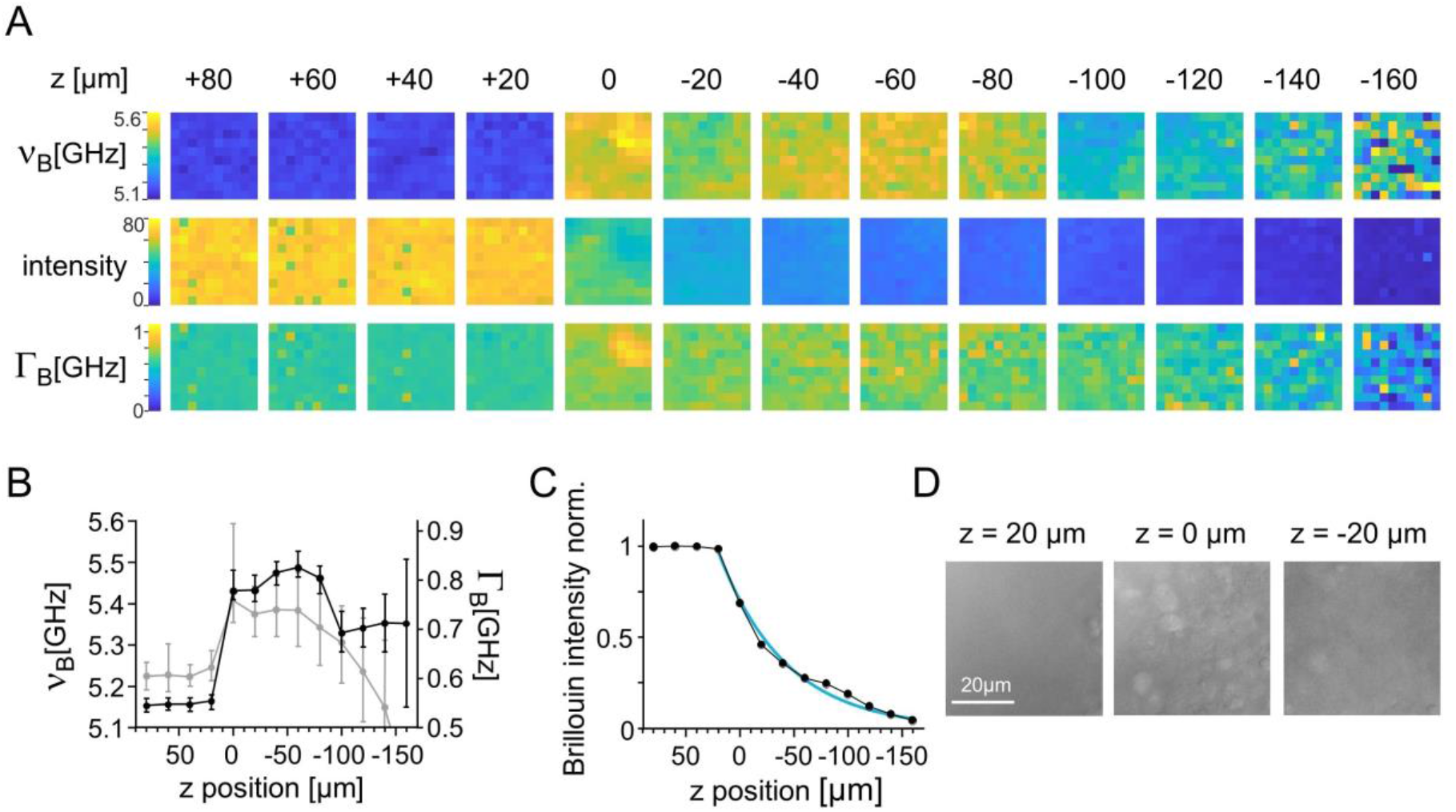
Effect of focal depth on Brillouin parameters of brain tumor tissue. **A:** Brillouin maps (10×10 points, 1 μm, step size) of a fresh tissue sample of squamous cell carcinoma brain metastases at various focal depths from z= 80 µm to z = -160 µm. **B:** Median Brillouin shift (black) and linewidth (gray) at different focal depths, error bars: 10% and 90% percentile. **C:** Median Brillouin intensity (normalized) in different focal depths. **D:** Bright field images of the measurement position at focal positions as indicated.

In deeper tissue layers, reduced median values for ν_B_ are accompanied by a substantial increase in the range values in the maps. The same phenomenon is found for Γ_B_. The reduced median values for ν_B_ and Γ_B_ can be attributed to the fitting algorithm, which employs initial values within this range. In the absence of sufficient signal, the algorithm is unable to converge to a local minimum, leading to values that are proximate to the initial values. The data indicates that valid Brillouin measurements can be performed up to a depth of z = -80 µm with this system and acquisition parameters. Low signal intensities at z = -100 µm and deeper are insufficient for accurate data analysis. The extraction of reasonable values for ν_B_ and Γ_B_ is strongly impacted because the fitting algorithms used might randomly assign peak positions leading to incorrect and widely distributed results as found for z = -160 µm.

Given the inherent heterogeneity of tissue and its uneven surface, factors other than the z-position might also affect the Brillouin signal intensity, especially when acquiring larger maps. Figure 3A shows an example of a Brillouin mapping of 100 ×100 µm on a GBM organoid. At position 1 (as labeled in Figure 3A, point 19,31 of the map), the Brillouin signal is intense and fitting procedures easily identify the Brillouin bands, allowing precise determination of ν_B_ and valid calculation of Γ_B_ (Figure 3B gray curve). However, inspection of the intensity map reveals several areas of very low intensity (Fig. 3A arrows). Those are 15-20 µm in diameter and might represent cells that are located above the focal position and attenuate the light. Problems with data fitting procedures can be identified by occurrence of random values within large ranges in the corresponding maps for ν_B_ and more pronounced in the map for Γ_B_. Examples of two Brillouin spectra acquired within the position 2 and the respective fitted bands are shown in Figure 3B. They are characterized by very low intensities (∼4) and fitting of points results in non-sense curves. Conclusively, the values for ν_B_ and Γ_B_ differ a lot for those adjacent points (position 2a (38,30) ν_B_ = 5.39 GHz and Γ_B_ = 1.31 GHz; position 2b (38,31) ν_B_ = 5.18 GHz and Γ_B_ = 0.67 GHz), and are not related to any tissue properties.

**Figure 3:**
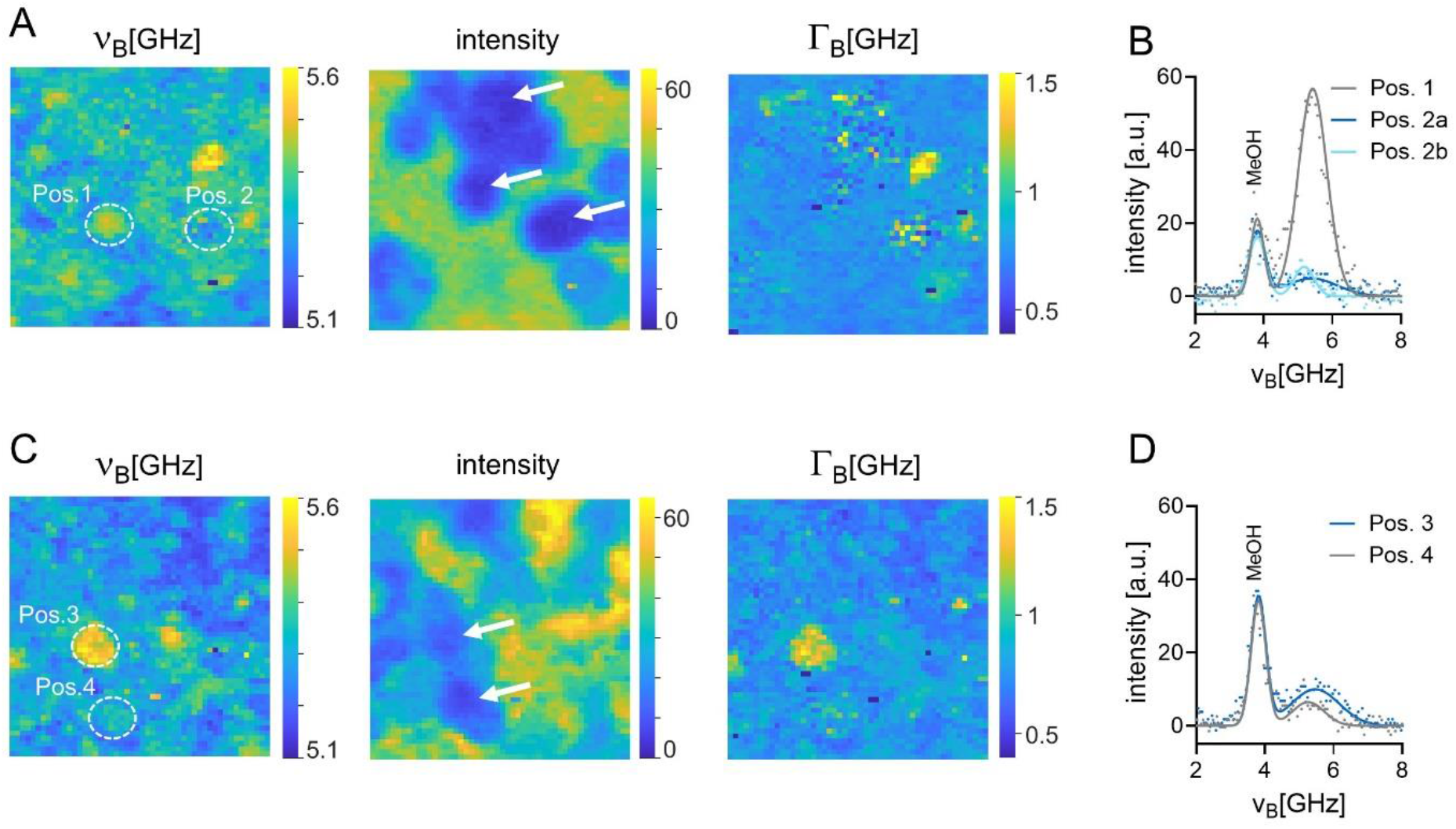
Effect of tissue structure on Brillouin shift parameters of brain tumor tissue. **A:** Brillouin maps acquired on a brain tumor organoid. **B:** Brillouin raw spectra (points) acquired in the areas indicated in A in the Brillouin shift map and corresponding fitted bands (lines). **C:** Brillouin maps of a fresh brain tumor sample (brain metastases of squamous cell carcinoma) **D:** Brillouin raw spectra (points) acquired in the areas indicated in C and corresponding fitted bands (lines) with bands of tissue and of reference methanol (MeOH) ν_B_ = 3.81 GHz. Arrows indicate areas with low Brillouin intensity. All maps: 50×50 points, 2 μm step size.

Figure 3C shows an example of Brillouin mapping of a fresh brain tumor sample (brain metastases of squamous cell carcinoma). In the Brillouin intensity map, structures above the focal plane can be identified (arrows) as in the previous example. However, the Brillouin intensity is slightly higher (values ∼10) and no random values were observed in the map for ν_B_ and Γ_B_. Neighboring pixels have similar values thereby reflecting the presence of tissue structures. The fitting procedure results in reasonable Brillouin bands (Figure 3D) and allows detection of differences in ν_B_ for positions 3 (point 15,26) and 4 (point 19,37). The data suggests that cells situated above the focal plane might affect the Brillouin signal in different ways. The presence of lipid-laden cells in GBM organoids might result in signal attenuation, while other types of brain tumor cells cause less signal loss. This underlines that consideration of regional variations in Brillouin intensity is important in analysis of Brillouin datasets.

Figure 4 shows Brillouin maps with 75 × 75 points and a size of 150 × 150 µm of fresh tumor samples. An uneven sample surface is a frequent problem when analyzing larger areas. The ν_B_ and intensity maps can be used to judge the focal position and evaluate the quality of the measurement. Figure 4A shows an example of a brain metastasis of lung cancer. The upper left and lower right corner show the ν_B_ ∼5.1-5.2 GHz and high Brillouin intensities and represent, therefore, measurements in the ACSF above the sample’s surface (Figure 4A asterisk). Moreover, small round structures with a diameter of 7-8 µm are visible in the ν_B_ map (Figure 4A) and the corresponding bright field image (Figure 4B), which is the size of human erythrocytes. The corresponding Raman spectrum (Figure 4C) shows the characteristic bands of hemoglobin at 750 and 1546 cm^-1^ [35] and confirms contaminations with blood in the field of view. Therefore, the Brillouin signals originate from the blood cells and do not represent the properties of the brain tumor sample. Figure 4D shows Brillouin maps of a case of meningioma and a small out of focus area (ACSF) was detected in the large maps (asterisk). The bright field image (Figure 4E) shows the tissue surface with a fiber-like structure, which can also be anticipated in the ν_B_ map. The Raman spectrum (Figure 4F) shows the characteristic bands of meningioma tissue [35,36] without blood contamination, confirming that the Brillouin data reflects the viscoelastic properties of the fresh brain tumor sample.

**Figure 4:**
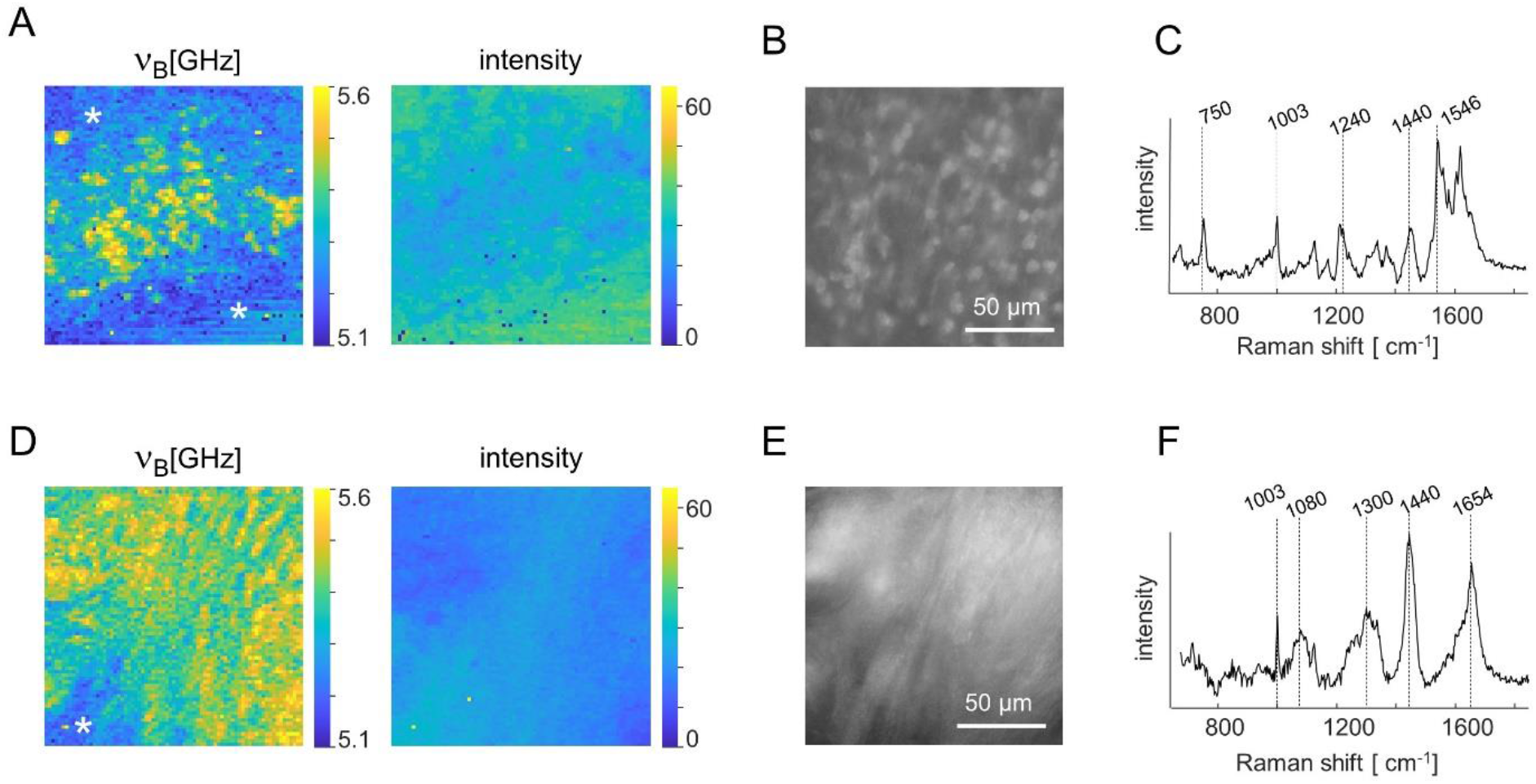
Effect of blood contaminations on Brillouin shift parameters of brain tumor tissue. **A:** Brillouin maps acquired on a fresh brain tumor sample (brain metastasis of lung cancer). **B/C:** Bright field image and Raman spectrum of the measurement position of A **D:** Brillouin maps acquired on a fresh brain tumor sample (meningothelial meningioma CNS WHO grade 1). **E/F:** Bright field image and Raman spectrum of the measurement position of D. Asterisks indicate out of focus areas. All maps: 75×75 points, 2 μm step size.

Figure 5 shows examples of colocalized Raman and Brillouin spectra obtained from two brain tumor organoids. Here, the analysis of the Raman spectral bands allows one to deduce the cellular compartment in which the measurement was performed. In the first example, the positions can be related to the cytoplasm (Fig. 5A, gray spectrum) or to the cell nucleus (Fig. 5A, blue spectrum) based on the Raman band at 782 cm^-1^ which is assigned to DNA [37]. The second example shows one position representing the cytoplasm (Fig. 5B, gray spectrum) and increased lipid-related bands at 1260, 1300 and 1440 cm^-1^ [37] at the second position (Fig. 5B, blue spectrum). In both cases, the cytoplasm exhibited a ν_B_ of 5.39 GHz. Higher values for ν_B_ of 5.48 GHz and 5.49 GHz were found for cell nuclei and lipid rich regions, respectively.

**Figure 5:**
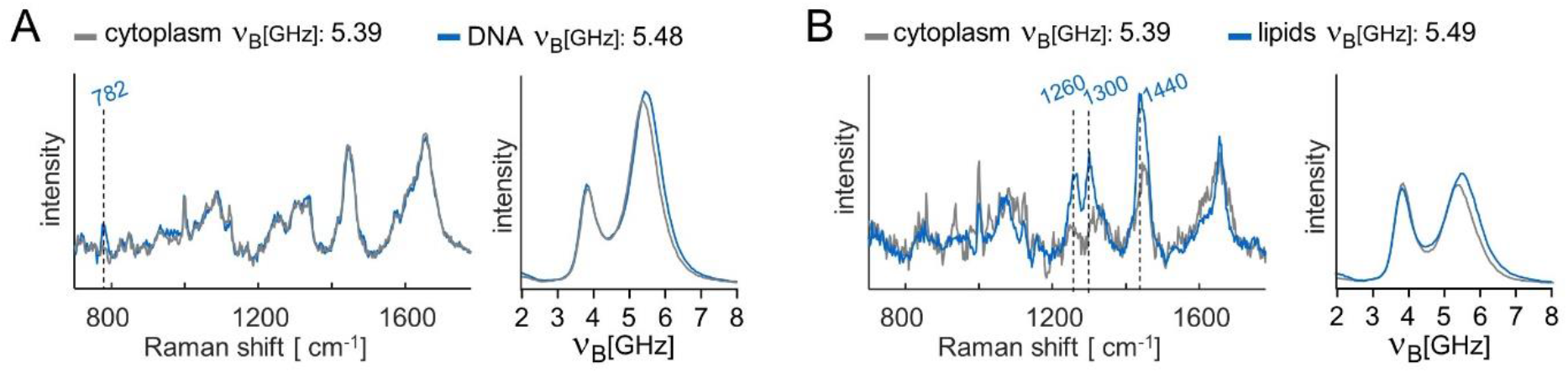
Brillouin shifts of different cellular compartments of brain tumors. **A/B:** Raman and Brillouin spectra acquired simultaneously from the same position of brain tumor organoids. Data from two positions on the same sample are shown. A and B represent measurement on different samples. Distance between positions: 5 µm and 20 µm for A and B, respectively.

## 4. Discussion

Performing Brillouin microscopy on brain tissue poses several challenges compared to the analysis of transparent samples such as water, gels or cornea. Nervous tissue exhibits attenuation properties that can vary significantly at the local level. Moreover, the complex three-dimensional architecture and intratumoral heterogeneity of human brain tumor samples, which results in highly localized variations in the Brillouin signal, need to be correctly interpreted in the tissue context. The data suggests that an increase in the Brillouin shift is specifically related to the biochemical signatures of DNA or lipids in brain tumors. For high lipid content, this might rather reflect changes in optical tissue properties than altered biomechanics [29]. These factors may also be relevant for discriminating between normal brain tissue and tumors. However, future systematic studies on large cohorts are required to address this issue.

We showed that consideration of tissue structure, spatial orientation as well as possible contaminations are of paramount importance for the analysis and interpretation of Brillouin data sets. While the Brillouin intensity does not carry information on viscoelastic properties of the sample, our data illustrate its indispensability for understanding Brillouin data quality, which might vary among different positions of the same map. In cases where the Brillouin intensity is too low, increasing the acquisition time or laser power may be an option. However, when working with live tissue, total measurement time and photodamage define certain limits for these parameters [18]. Working in the NIR range like in this study constitutes an advantage because of higher penetration depth and potentially less phototoxicity [38].

With our system, valid data sets can be obtained up to a depth of ∼80 µm in fresh nervous tissue. This is comparable to data of other groups who successfully performed Brillouin mapping on mouse brain 50 µm beneath the surface [25] and in a depth of ∼90 µm and ∼100 µm on ovarian cancer nodules [39] and chicken muscle [40], respectively. In more transparent tissue, larger penetration depth up to some hundred micrometers were found on mouse [41] and zebrafish embryo [34]. This underpins that the penetration depth of Brillouin microscopy has to be evaluated for each system and experimental series as it depends on excitation, measurement geometry and tissue type. In our study, this means that Brillouin microscopy is able to address cells in some layers below the surface given the typical diameter of cells of the nervous system of 10-20 µm. Those cells might be less impaired by cutting and slicing procedures than those of the first cell layer.

We demonstrated that presence of certain cell types and blood contamination of the surface of the tissue of interest strongly impair the measurement. For future ex vivo studies on human brain tissue, additional washing steps might help to avoid blood contamination and visual inspection of bright field microscopic images of the sample surface could help to find areas without erythrocytes if no other ancillary technique such as Raman spectroscopy is available. In addition to providing a reference for contamination, Raman spectroscopy can offer biochemical data sets to aid in the interpretation of Brillouin spectra [42,43]. Both technologies operate optically and can share laser excitation resulting in colocalized, simultaneous measurements [44].

While comprehension and correct handling of artefacts is of paramount importance for meaningful measurements, retrieving tissue longitudinal modulus poses further challenges. A conversion of the Brillouin frequency shift to the longitudinal elastic modulus was not performed in the present study, as knowledge of the local mass density and refractive index is necessary for this, which is the main limitation of the technique. As brain tissue is locally heterogeneous, measurements of both quantities co-registered with the Brillouin measurement are required to enable the conversion of the Brillouin shift to the real part of the longitudinal modulus. Such measurements require special instrumentation. In biological applications, analysis is typically focused on relative variations in the Brillouin shift. Changes in longitudinal modulus are inferred by assuming that the Lorentz-Lorenz relation is valid, meaning that the variations in the squared refractive index and mass density cancel each other out in a heterogeneous sample [45,46].

Local tissue hydration and optical scattering phenomena within the tissue can further complicate data interpretation in terms of tissue viscoelastic properties. All measurements were performed in solution (ACSF, medium), leading to an equilibrium and mimicking in vivo situation. Nevertheless, hydration constitutes a significant issue. Tissue and cells can be described as composed by a solid solute (the cell material) in water, where tissue compressibility is the sum of compressibility of water and solid part weighted on their respective volume fractions [47,48]. Therefore, the Brillouin shift is also sensitive to changes of tissue water content and the relative changes of Brillouin shift are about tenfold more sensitive to changes in water content than to changes in the compressibility of the solid part [47,49]. Therefore, caution is required by interpreting the Brillouin shift as proxy for the elastic properties of the solid part of cells and tissue, as the local solute concentration changes within nervous cells and tissue [50,51]. Also in this context, Raman spectroscopy can help in the interpretation [52].

The Brillouin linewidth depends from viscosity, but heterogenous broadening [53] and multiple scattering within the tissue [54] lead to further spectral broadening in Brillouin spectroscopy of biological samples. Moreover, system-related broadening becomes important when using high numerical aperture objectives [55]. The system broadening can be measured and in our experiments accounts for 0.33 GHz. On the other hand, heterogeneous broadening and multiple scattering depend from the local tissue characteristics and must be eventually evaluated on each dataset. Strong multiple scattering was reported to alter the symmetry of the bands in the low frequency region, as shown with green excitation and a high contrast, high spectral resolution spectrometer at depths of some hundreds of micrometers in turbid medium [54]. We could not observe this effect in our Brillouin spectra, also because our system can provide much lower contrast and resolution, and the MeOH bands partly obscure the low frequency region. Moreover, this effect might be less prominent as we used a near infrared excitation to reduce the scattering.

The combination of Brillouin datasets with datasets from other mechanical techniques is possible and may be used for better interpretation of Brillouin measurements. Multimodal mechanical mapping has been proposed to provide comprehensive insight into the biomechanical properties of tissues [56], which may help to understand malignant transformation and mechanisms of invasion mechanisms in cancer [30]. OCT can be combined with Brillouin microscopy [57] to provide guidance and reveal tissue micromorphology [58]. Towards intraoperative use, OCT might be used for motion correction during Brillouin microscopy [59]. Furthermore, OCE can be applied to calibrate Brillouin measurements and retrieve the sample-dependent relationship between the Brillouin-deduced longitudinal modulus and Young’s modulus [60]. It has the potential to be applied intraoperatively using an air jet to apply the mechanical load [61]. Indentation experiments can provide additional information on stress relaxation, which might help distinguishing tumorous from non-tumorous tissue [9].

Technically, Brillouin spectroscopy is a challenging technique. However, as non-contact and label-free technology it has few barriers to clinical translation and initial in vivo measurements have been demonstrated in animal models [34,62,63], on the crystalline lens [64] and on human skin [65]. Moreover, recent research demonstrated fast imaging using line scan and full-field Brillouin microscopy, thus reducing phototoxicity and allowing monitoring of dynamic processes [66–68]. However, several technical hurdles which are well recognized in the broader context of in vivo optical imaging have to be considered also for Brillouin microscopy in neurooncology. This includes motion artifacts particularly from respiration and cardiac cycles, which can affect the spatial and spectral resolution of Brillouin measurements, and environmental fluctuations such as temperature variations, which may alter refractive index or shift the Brillouin peak position. Moreover, tissue relaxation and mechanical adaptation after craniotomy and during longer acquisition times could influence the mechanical state of the tissue during imaging. Therefore, routine in situ Brillouin microscopy will require significant additional technical developments, particularly in stabilization, real-time processing, and signal correction algorithms. Obviously, it would be particularly important for future clinical applications to recognize the limitations of the technology and the measurement situation and to define safe corridors for obtaining reliable and meaningful data sets.

## Declarations Ethical Statement

All patients provided written informed consent and the study was approved by the ethics committee of TU Dresden (EK 323122008, date: 17.02.2009).

## Data availability statement

The datasets generated during the current study are available from the corresponding author on reasonable request.

## Acknowledgements

None

**Acknowledgements**

The authors thank the National Center for Tumor Diseases (NCT), Partner Site Dresden, for supporting the experimental set-up.

## Author contributions:CRediT

Conceptualization: OU, RG

Formal Analysis: OU, RG

Investigation: TL

Methodology: JR, RG

Resources: AT, IYE

Software: JR

Supervision: OU, RG

Visualization: TL, OU

Writing – original draft: OU, RG

Writing – review and editing: all authors

